# FICD deficiency protects mice from hypertrophy-induced heart failure via BiP-mediated activation of the UPR^ER^ and ER-phagy

**DOI:** 10.1101/2024.05.28.596287

**Authors:** Shannon M. Lacy, Rebecca J. Taubitz, Nicholas D. Urban, Samantha N. Turowski, Eric D. Smith, Adam S. Helms, Daniel E. Michele, Matthias C. Truttmann

**Author notes:** To whom correspondence should be addressed: BSRB, 109 Zina Pitcher Place, Ann Arbor 48109, MI.

## Abstract

Cardiomyocytes require the HSP70 chaperone BiP to maintain proteostasis in the endoplasmic reticulum (ER) following cardiac stress. The adenylyl transferase (AMPylase) FICD is increasingly recognized to regulate BiP activity through the post-translational addition of an adenosine monophosphate moiety to BiP surface residues. However, the physiological impact of FICD-mediated BiP regulation in the context of cardiovascular health is unknown. Here, we find that FICD deficiency prevents pressure overload-associated heart failure, hypertrophy, and fibrosis, and that FICD knockout mice maintain normal cardiac function after cardiac pressure overload. At a cellular level, we observe that FICD-mediated BiP AMPylation blunts the induction of the unfolded protein response (UPR^ER^) and impairs BiP interaction with FAM134B, an ER-phagy receptor, thus limiting ER-phagy induction under stress. In contrast, FICD loss significantly increases BiP-dependent UPR^ER^ induction and ER-phagy in stressed cardiomyocytes. We also uncover cell type-specific consequences of FICD activity in response to ER stress, positioning FICD as a critical proteostasis regulator in cardiac tissue. Our results highlight a novel regulatory paradigm controlling stress resilience in cardiomyocytes and offer a rationale to consider FICD as a therapeutic target to treat cardiac hypertrophy.

## Introduction

Cardiac hypertrophy is a leading cause of heart failure^1–3^. Risk factors for this cardiovascular condition include advanced age, obesity, diabetes, and high blood pressure^4–7^. A hallmark of cardiac hypertrophy is the accumulation of misfolded proteins and a decline in cardiac protein homeostasis, or proteostasis^8–11^. However, the characterizations of the mechanisms that regulate cardiac proteostasis is limited.

Cardiomyocytes and cardiac fibroblasts are among the most abundant cells in the heart and are central to proper functioning of cardiac muscle^12–14^. Cardiomyocytes are terminally differentiated, long-lived cells that depend on efficient protein folding and recycling machinery to maintain contractile function and cellular survival^15–17^. Cardiac fibroblasts are secretory cells essential to maintain cardiac structural integrity and the extracellular matrix^18,19^. In both cardiomyocytes and cardiac fibroblasts, the endoplasmic reticulum (ER) is the central location of protein synthesis and regulation of proper protein folding^20,21^. During cardiac stress induced by pressure-overload, unfolded/misfolded proteins accumulate in the ER and challenge proteostasis^22,23^. To restore proteostasis in the ER, stressed cardiomyocytes and cardiac fibroblasts induce the Unfolded Protein Response (UPR^ER^) as well as ER-selective autophagy (ER-phagy), thus mitigating the accumulation of misfolded proteins and preventing ER-dependent apoptosis^24–27^.

Binding Immunoglobulin Protein (BiP; Grp78) is an essential ER resident chaperone involved in protein folding as well as the regulation of the UPR^ER^ and autophagy during ER stress in the heart^28–30^. In stress-free conditions, BiP complexes with and thus inactivates UPR^ER^ sensors, IRE1α, PERK, and ATF-6, in the ER lumen^31–33^. The accumulation of misfolded proteins in the ER promotes the dissociation of BiP from these sensors, which enables the subsequent induction of UPR^ER^-mediated transcriptional changes ^34–36^. IRE1α and PERK are kinases that dimerize and autophosphorylate upon BiP dissociation^37,38^. Dimeric PERK phosphorylates Elf2α, resulting in a reduction of protein translation while upregulating autophagy- and lysosome-related genes^39–41^. Dimeric IRE1 cleaves full length X-box binding protein 1 (XBP1) mRNA, producing the transcription factor, spliced XBP1, that translocates to the nucleus and binds to downstream UPR gene promoters^42,43^. The ATF6 receptor translocates to the Golgi upon activation, where it is cleaved and subsequently enters the nucleus to bind to UPR-related gene promoters^43^. Upon excessive ER stress, the three pathways trigger the transcription of the pro-apoptotic factor, CHOP, linking excessive ER stress to increased cellular death^44,45^.

In cardiomyocytes, autophagy is critically involved in the recycling of cellular material encapsulated in autophagosomal vesicles via the binding to and degradation by the lysosome^46–48^. While lysosomal fusion and degradation occur in the cytosol, the initiation of autophagy can occur in various cellular compartments^49,50^. One such process is ER-phagy, referring to the selective degradation of parts of the ER through engulfment by a phagophore membrane^51,52^. This process is mediated through ER-phagy receptors in the ER membrane which bind to LC3 on the phagophore membranes^51,52^. BiP regulates ER-phagy through at least two mechanisms: First, BiP mediates PERK activation and thus the expression of genes required for autophagic protein degradation^40,53,54^. Second, BiP directly binds to the luminal domain of the ER-phagy receptor, FAM134B, which initiates ER-phagy during ER stress^55^. Despite the critical role of BiP in maintaining ER proteostasis in cardiomyocytes, our understanding of the mechanisms that regulate BiP activity remains incomplete.

FICD is a mammalian enzyme that confers the post-translational modification (PTM), AMPylation, to substrate proteins^56–60^. Using a single catalytic site, FICD both adds (AMPylation) and removes (deAMPylation) adenosine monophosphate (AMP) to threonine and serine residues on protein substrates^61–64^. The best characterized FICD substrate is BiP, which can be AMPylated on at least two residues (Thr-366, Thr-518)^57,58,63,65–70^. FICD-mediated AMPylation locks BiP in an adenosine triphosphate (ATP)-bound conformation unable to hydrolyze ATP or interact with client proteins^63,67,71,72^. Previous studies in non-cardiac cells indicate that both excessive and reduced AMPylation can lead to ER-dependent apoptosis and limit UPR^ER^ induction under stress. However, the direct impact of BiP AMPylation on UPR activity *in vivo* remains poorly understood^58,68,73,74^.

BiP is required for maintaining proper cardiac contractility and function^75,76^. While BiP deficiency in cardiomyocytes is embryonic lethal, BiP ablation in the adult cardiac muscle leads to increased apoptotic cell death and a decline in heart performance^30^. Understanding how FICD activity regulates BiP in cardiomyocytes and other cell types may provide novel insights into mechanisms regulating cardiac stress resilience and protein unfolding stress during pathological cardiac remodeling.

In this study, we use a transverse aortic constriction (TAC) model of cardiac pressure overload to show that FICD deficiency preserves cardiac ejection fraction (EF) and protects the heart from pressure-induced hypertrophy and fibrosis. Utilizing primary cardiomyocytes and cardiac fibroblasts, we find that these benefits are mediated through increased activation of the UPR^ER^ and ER-phagy. We further demonstrate that the consequences of FICD activity in the heart are cell type-specific, as cardiac fibroblasts and cardiomyocytes show distinct non-overlapping patterns of BiP AMPylation and FICD-dependent CHOP activation in responses to ER stress. Finally, we show that FICD deficiency potentiates UPR^ER^ induction and BiP-dependent ER-phagy in stressed primary cardiomyocytes. Taken as a whole, our results suggest that FICD deficiency enhances cardiomyocyte and cardiac fibroblast stress resilience through the derepression of UPR^ER^ signaling and ER-phagy, leading to increased functional capacity in stressed hearts.

## Materials and Methods

### Antibodies

The following antibodies were used for western blotting and immunofluorescence studies: Alpha-actinin (D6F6)(Cell Signaling, 6487T, Rabbit mAb), BiP (Proteintech, 665741IG, mouse mAb), CHOP (D46F1)(Cell Signaling, 5554S, Rabbit mAb), Cathepsin B (D1C7Y)(Cell Signaling, 31718S, Rabbit mAb), Cathepsin D (E179)(Cell Signaling, 69854S, Rabbit mAb), Cathepsin L (Santa Cruz Biotechnology, sc-32320, mouse mAb), FAM134B (E8Y9R)(Cell Signaling, 83414S, Rabbit mAb), GAPDH (Proteintech, 600041IG, mouse mAb), HSC70 (B-6)(Santa Cruz Biotechnology, sc-7298, mAb), LAMP1 (E6N3R)(Cell Signaling, 46843S, Rabbit mAb), LC3A/B (D3U4C)(Cell Signaling, 12741S, Rabbit mAb), Thr-AMPylation (Biointron, 17G6, mAb), Secondary 647 Fluorescent antibody (ThermoFisher, A32728).

### Animal Models

All animal procedures were performed at the University of Michigan facilities according to NIH guidelines and approved by the Animal Care and Use Program (ACUO), the Unit for Laboratory Animal Medicine (ULAM), and the Institutional Animal Care and Use Committee (IACUC) guidelines at the University of Michigan. C57BL/6 purchased from The Jackson Laboratory and full body *Ficd^-/-^* mice ^77^ were used for this study. All mice were fed a standard chow diet and water ad libitum, and housed in ventilated cages with a 12-hour light and dark cycle.

### TAC and Sham Experiments

TAC and sham surgery were performed according to previously described methods ^78^. 6-month or 12-month mice were given carprofen pre-emptively and for 48 hours post-surgery, then as needed for analgesia. The mice were anesthetized with isoflurane and orally intubated and ventilated. The skin over the ventral surface of the neck was incised with scissors from below the chin caudally to above the xyphoid. The mandibular gland was retracted cranially. A small, curved forceps was used to carefully enter the extra-pleural space under the manubrium. The tissue along the sternum was cauterized over the manubrium and caudally to the third sternal plate. The sternum from the manubrium to the third sternal plate was incised down the middle. The thoracic opening was retracted left and right with 4-0 braided non-absorbable suture. The aortic arch was isolated by entering the extra-pleural space above the first rib, and the transverse aorta was isolated between the right and left carotid arteries. For the TAC surgery, a 6-0 size silk suture ligature was tied around the transverse aorta against a 27G needle to produce a constriction after the removal of the needle. For the sham surgery, the aorta was manipulated in the same manner as in the TAC surgery, without placing a ligature around the aorta. The thoracic opening was closed with the retractor sutures. The incision was closed with three stainless steel wound clips and tissue adhesive was applied to reinforce the close. The mice recovered on heating pads after surgery and were subsequently returned to their cages. 8-weeks after the surgeries, when the cohorts were 8-months or 14-months-old, respectively, the hearts were collected for biochemical and histological analysis, using liquid N2 snap freezing for protein and RNA analysis and liquid Nitrogen cooled-isopentane freezing in OCT for histological analysis.

### Echocardiographic Assessment of Cardiac Function

6-month and 12-month animals underwent echocardiograph assessment 2 weeks before, 2-weeks after, 4-weeks after, and 7-weeks after TAC or sham surgery. Echocardiographic assessments were performed as previously described ^78–80^. Induction of anesthesia was performed in an enclosed container filled with 3-5% isoflurane. After induction, the mice were placed on a warming pad to maintain body temperature. 1-1.25% isoflurane was supplied via a nose cone to maintain a surgical plane of anesthesia. The hair was removed from the thoracic area with depilatory cream. The mouse was placed on a heated platform to prevent hypothermia and ECG and respirations were monitored via non-invasive resting ECG electrodes. Eye lubricant was applied to prevent corneal damage during prolonged anesthesia. Transthoracic echocardiography was performed in the supine or left lateral position. Two-dimensional, M-mode, Doppler and tissue Doppler echocardiographic images were recorded using a Visual Sonics’ Vevo 2100 high resolution in vivo micro-imaging system with a MS 550D transducer which has a center frequency of 40 MHz and a bandwidth of 22-55 MHz and a MS250S transducer for the TAC animals with a bandwidth of 13-24MHz. We measured LV ejection fraction from the two-dimensional long axis view. We measured systolic and diastolic dimensions and wall thickness by M-mode in the parasternal short axis view at the level of the papillary muscles. Fractional shortening and ejection fraction were also calculated from the M-mode parasternal short axis view. Diastolic function is assessed by conventional pulsed-wave spectral Doppler analysis of mitral valve inflow patterns (early [E] and late [A] filling waves). Doppler tissue imaging (DTI) is used to measure the early (Ea) diastolic tissue velocities of the septal/and lateral annuli of the mitral valve in the apical 4-chamber view.

### Immunohistochemistry (Fibrosis and H&E)

Frozen mouse hearts in OCT were sectioned onto glass slides using a Leica CM3030S Cryostat and stored frozen until ready to be stained. Hematoxylin and eosin staining (H&E) and Picro-Sirius red staining was performed using rehydrated samples following standard protocols. Briefly, frozen slide sections were brought to room temperature for five minutes and air dried. The sections were fixed using 10% buffered formalin for 5 minutes and rinsed with running water for 5 minutes. For H&E staining, the sections were stained with Hematoxylin (Sigma-Aldrich, MHS1) and Eosin (Sigma-Aldrich, HT110132). For fibrosis staining, sections were stained with Sirius Red stain, made from Picric acid (Sigma-Aldrich, 197378), Direct Red (Sigma-Aldrich, 365548), and Fast Green (Sigma-Aldrich, F7252). The slides were rinsed serially with water, dehydrated serially with Ethanol, and cleared serially with Xylenes. Tissues were mounted with glass coverslips using Permount (Electron Microscopy Sciences, 1798601) and dried overnight. Slides were scanned at 20x and stitched together using the Keyence BZ-X Viewer microscope and BZ-X Analyzer software. For cardiomyocyte hypertrophy quantification, 25 cells from 5 sections of the same left ventricle heart were outlined in ImageJ and the Feret’s minimum diameter was measured. For cardiac fibrosis quantification, sections from three depths of the same heart ventricle were analyzed using ColorDeconvolution2 in ImageJ to separate the stained-red fibrotic tissue from other stained-green cardiac tissues, and the percent fibrosis was calculated.

### Cell Culture and Transfection

Neonatal primary cardiomyocytes and cardiac fibroblasts were isolated from WT and *Ficd^-/-^* mice (post natal day 1-3) as previously described ^81^. Briefly, 6-8 hearts were washed in 20mM butanedione monoxime (BDM)(Sigma-Aldrich, B-0753) in 1x phosphate buffered saline (PBS) on ice, transferred to isolation buffer (20mM BDM, 0.0123% trypsin in Hank’s balanced salt solution without Ca^2+^, Mg^2+^), and minced with sterile scissors. The minced hearts were incubated overnight at 4°C in isolation buffer with gentle agitation. The minced hearts were transferred to digestion buffer (20mM BDM, 1.5mg/ml Roche Collagenase/Dispase enzyme mix in Leibovitz L15 medium)(Roche, 10269638001)(cellgro, 10-045-CV) and agitated at 60rpm at 37°C for 25 minutes. The cells were further dissociated by resuspending the mixture 20 times in a serological pipette and then strained through 100μm cell strainer. Cardiomyocytes were centrifuged at 500rpm for 10 minutes at 20°C, and cells were resuspended in plating media (65% DMEM high glucose, 19% M-199, 10% horse serum, 5% fetal calf serum, 1% penicillin/streptomycin)(cellgro, 10-013-CV)(10-060-CV)(cellgro, 35-030-CV)(35-011-CV)(cellgro, 30-002,CI) and plated into 2 wells of a 6 well plate. Fibroblasts adhered to the non-coated 6 well plate for 90 minutes at 37°C. Non-adherent cardiomyocytes were washed from the plate and re-plated in 2 wells of gelatin and collagen pre-coated 6-well plates. Plating media was re-added to the cardiac fibroblasts that had adhered to the first non-coated plate. Isolated cells were maintained in plating medium for 24 hours. Transfections were performed 24 hours after initial plating in low serum media. Media was replaced 24 hours after transfection. For ER stress assays, cells were treated for 12 hours with either Tunicamycin (10μg/mL) or Thapsigargin (3μM) before harvesting for protein or mRNA analysis. For autophagy and ER-phagy flux assays, cells were treated with bafilomycin A (100nM)(Sigma-Aldrich, SML1661) for 4 hours prior to harvesting for protein analysis. For assays requiring increased levels of AMPylation, the plasmid encoding a constitutively active FICD mutant (E234G) in a pcDNA3.1 backbone was used.

### Cardiomyocyte Contractility Analysis

The contractility function assessments of neonatal cardiomyocytes were performed using the two-dimensional cardiac muscle bundle (2DMB) technique, as previously reported^82^. Briefly, polydimethylsiloxane (PDMS) substrates were fabricated at a physiologic elasticity (8 kPa). Each PMDS substrate was micropatterned with fibronectin in an array of 7:1 aspect ratio rectangles (each 308μm x 45μm) to enable coordinated tissue formation within each rectangle. Cardiomyocytes were isolated as described previously from 6-8 mice, and 60,000 cardiomyocytes were plated in one mL of plating media in the inner well of each 2DMB plate for 1 hour at 37°C to allow attachment. Then, 4 mL of plating media was exchanged every day until contractile assay, which was performed three days following plating. Live cell microscopy was performed at 37°C and 5% CO2 using a Nikon Ti2 Widefield microscope with a 40x air objective. Cardiomyocyte muscle bundles that were contracting uniformly in individual rectanglular micropatterns were selected for contractile imaging. Contractile function was measured using ContractQuant, a previously published MATLAB method (available on GitHub: https://github.com/Cardiomyocyte-Imaging-Analysis)^82^ developed for tracking cardiomyocyte bundles. Regions of interest were placed equidistant across the center of the contracting tissue and the contractile velocity and fractional shortening values were assessed.

### Immunofluorescence

Isolated cardiomyocytes and cardiac fibroblasts were fixed for 10 minutes with 4% paraformaldehyde, washed with PBS and stored at 4°C. Cells were permeabilized with 0.1% Triton/PBS for 30 minutes, blocked with 5% Bovine Serum Albumin (BSA), and stained for one hour at RT with alpha-actinin to visualize sarcomere structure. Stained samples were washed and stored at 4°C in PBS, or mounted using Prolong^Tm^ Antifade Mountant (ThermoFisher, P36961) until imaging. Images were collected with the Nikon Ti2 Widefield Microscope microscope at 20x.

### Transmission Electron Microscopy

Cardiomyocytes were digested and isolated as described in the methods, and plated on fibronectin precoated glass bottom plates. Cells were washed three times with 1X PBS and fixed with 3% Glutaraldehyde (GA)/3% Paraformaldehyde (PFA). Fixed cells were stored at 4°C until post-fixation processing. Fixed cardiomyocytes were next washed with 0.1M Sodium cacodylate buffer (CB) three times for five minutes each, post-fixed with 1% K4Fe(CN)6 + 1% OsO4 in 0.1M CB for 15 minutes, washed with 0.1M CB three times for three minutes each, washed with 0.1M Na2+Acetate Buffer, pH 5.2 for three times for three minutes each, and underwent en bloc staining with 2% uranyl acetate (UA) in 0.1M Sodium Acetate Buffer, pH 5.2 for 15 minutes. Cardiomyocytes were then washed with ultra-pure water for 3 minutes and underwent serial ethanol and Epon dilutions for dehydration and polymerization into blocks for sectioning. The blocks were embedded for 30 minutes at RT and polymerized for 24 hours at 65°C, which was repeated twice. The blocks were sectioned using a Leica UC7 ultramicrotome at 70nm and samples were imaged on a JEOL JEM-1400plus transmission electron microscope at 60kv.

### Real Time Quantitative PCR

RNA was extracted from isolated cardiomyocytes from 6-8 mice (prepared as described above in 6 well plates), isolated cardiac fibroblasts from 6-8 mice (prepared as described above in 6 well plates), or from frozen total heart ventricle tissue. RNA extraction was performed with the Zymo Direct-zol^TM^ RNA MiniPrep Plus kit (Cat# R2070). cDNA was synthesized from purified RNA using the High Capacity cDNA Reverse Transcription kit (Cat# 4368814) from ThermoFisher. Real time quantitative PCR was performed with primers GCTGAGTCCGCAGCAGG and CAGGGTCCAACTTGTCCAGAAT (spliced XBP1), GGTGGATTTGGAAGAAGAGAACC and CATAGTCTGAGTGCTGCGGA (total XBP1), GCGCTGCGGTAGGATCA and GATTTCGTGAAGAGCGCCAT (ATF4), GAGGTGTCTGTTTCGGGGAA and GCAAACAACGTCGACTCCCA (ATF6alpha), TGTGTGTGAGACCAGAACCG and AACACACCGACGCAGGAATA (BiP), as well as TCCAGTAAAGAAGACACCCAGA and GACAAAACCAGCGAGACACA (EEF-1; internal control). The expression of mRNA from these targets was determined by real time quantitative PCR using a QuantStudio Real-Time PCR system with the cycle parameters as: 50°C 2 min, 95°C 10 min, 95°C for 15 sec and 60°C for 1 min which was repeated 40 cycles. Results were quantified using the 2^-DDCT^ equation from the CT-values from three replicates per condition.

### Western Blotting

Isolated cardiomyocytes from 6-8 mice per condition (plated and treated as described above) were lysed with RIPA-containing protease inhibitor buffer for 15 minutes on ice and sonicated at 20% for 30 seconds on ice. Cell debris was spun down for 15 minutes at 4°C, and protein quantification was performed used the DC^TM^ Protein Assay kit (Cat# 5000112) from BioRad. 10μg of cell lysate was run using 10% or 15% Tris-Glycine SDS acrylamide gels. Gels were run for 1.5 hours at 100V, and transferred to ethanol-activated 0.2μm PVDF membranes using the Trans-Blot Turbo RTA Mini Transfer kit (#1704272) from BioRad. Each blot was blocked using 5%/0.1% BSA/Tween in TBS and incubated with primary antibody overnight at 4°C. The blots were washed with 0.1% Tween/PBS three times, and probed with secondary HRP-conjugated antibody for one hour at RT. The blots were washed with 0.1% Tween/PBS three times and images were acquired using the ImageQuant^TM^ LAS 4000 biomolecular imager. Western band quantification was performed using the online Image Lab software from BioRad from three replicates per condition.

### Immunoprecipitation

Isolated cardiomyocytes were plated and treated as described in the methods, as well as lysed and quantified for protein concentration as described above. Cell lysates were pre-cleared using Pierce^TM^ Protein A Magnetic beads (#88846) from ThermoFisher for 1 hour at 4°C, and the beads were removed using a magnetic rack. 150μg of cell lysate was incubated with 0.6μg of FAM134B antibody overnight at 4°C. 50uL of Protein A Magnetic beads were added to the mixture for 2 hours at 4°C. The immunoprecipitated proteins were isolated from the cell lysate mixture using the magnetic rack. The isolated beads were washed three times with the cell lysis buffer. The bound proteins were released from the beads by adding SDS loading buffer and boiled for 10 minutes. These proteins were then run on 10% Tris-glycine SDS acrylamide gels along with 10% input samples and transferred onto PVDF membranes overnight at 4°C. The membranes were probed for FAM134B and associated proteins. The IP was performed three separate times and the bands were quantified using the online Image Lab software from BioRad.

### Cathepsin L Enzyme Activity Assay

The activity of Cathepsin L from isolated cardiomyocytes from 6-8 mice (plated as described) was assessed using the Cathepsin L Activity Assay kit from Abcam (ab65306). 50μg of protein from isolated cardiomyocytes were used to assess the degradation of the FR-AFC substrate at 37°C degrees for 2 hours, reflective of Cathepsin L *in vitro* activity. A BioTek Synergy 2 Microplate Reader was used to assess the amount of cleaved AFC (amino-4-trifluoromethyl coumarin) released from active Cathepsin L. The Cathepsin L inhibitor (CL Inhibitor) from the Abcam kit was added to the mixture as a negative control.

### Statistical Analysis

Statistical Analysis was performed using the GraphPad Prism (version 10.2.2) software. Unpaired or paired t-tests, 2-way ANOVA tests, and 3-way ANOVA tests were performed. Figure legends specify the utilized tests for each data panel. A p value of p <0.05 was used to determine statistical significance.

## Results

### FICD deficient mice are protected from hypertrophy-induced heart failure

Previous work by several labs established a critical role for FICD in mitigating protein misfolding stress in the ER, particularly in long-lived terminally differentiated non-cardiac cells^83,84^. We hypothesized that FICD regulates ER proteostasis in cardiomyocytes, which experience protein misfolding stress during cardiac hypertrophy. To test this, we performed sham and TAC surgery on six-month-old C57BL/6 wildtype (WT) and FICD knockout (*Ficd^-/-^)* mice and assessed cardiac function using echocardiography at two-, four-, and seven-weeks post-surgery (Fig. 1A). Pressure gradient measurements across the TAC ligatures on the aortas confirmed that the procedure induced comparable pressure overloads in all hearts (Supplemental Fig. S1A). As expected, the ejection fraction (EF) of 6-month-old WT mice had significantly declined seven weeks post-TAC surgery compared to sham controls, indicating a decline in heart performance (Fig. 1B). In contrast, age-matched *Ficd^-/-^* mice maintained EF levels after TAC surgery at all tested time points (Fig. 1B). Two-way ANOVA analysis confirmed a significant contribution of the genotype on the consequences of TAC surgery on EF (P_int_ = 0.003) (Supplemental Fig. S1B).

**Figure 1.**
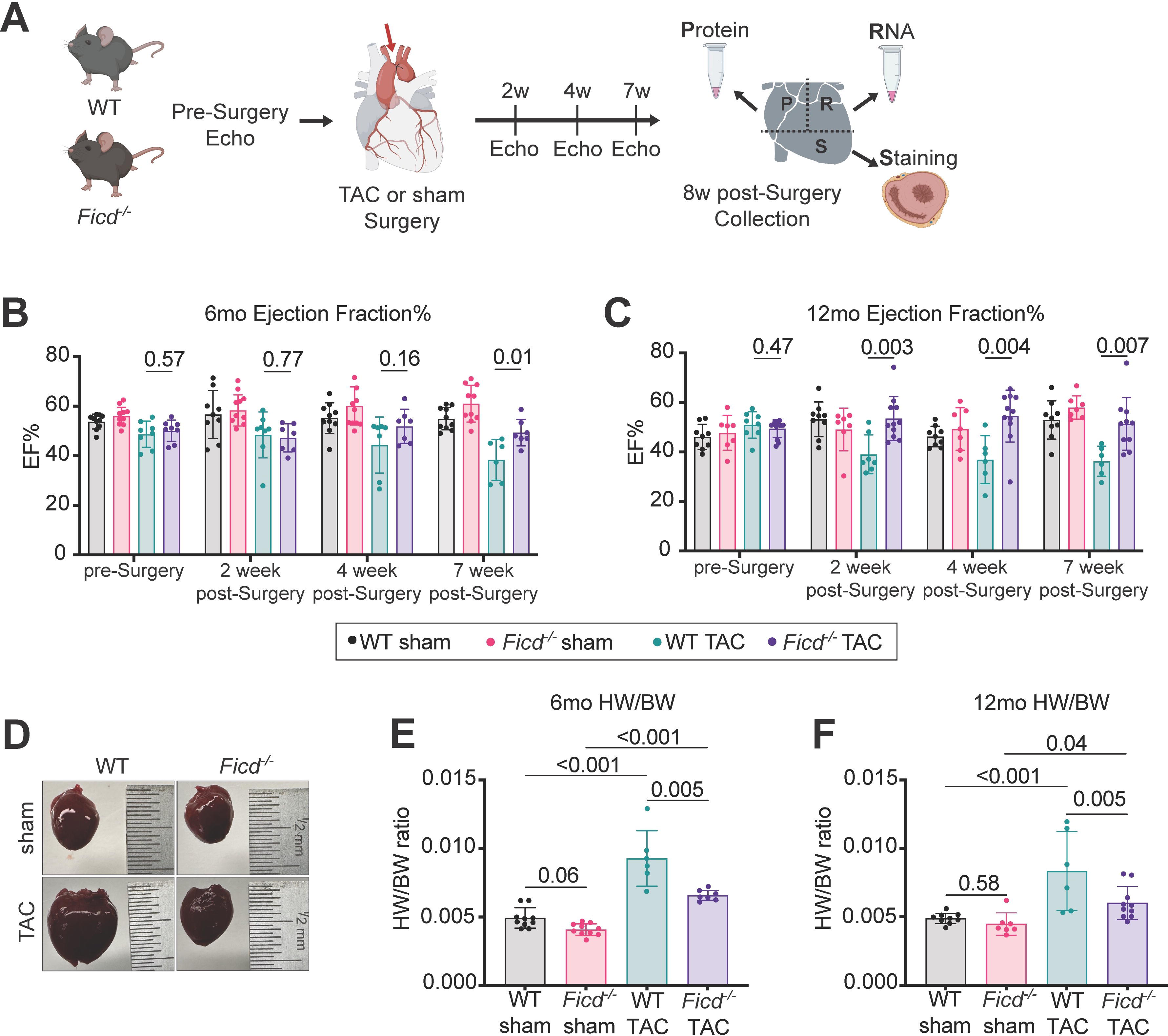
FICD deficient mice are protected from hypertrophic-induced heart failure. (A) Schematic of the experimental design to assess mouse heart function, structure, and associated molecular changes. (B) EF echocardiographic results from 2D long axis imaging of 6-month-old mice (n= 8-10 per genotype per surgery) after sham or TAC surgery. (C) EF echocardiographic results from 2D long axis imaging of 12-month mice (n= 7-12 per genotype per surgery) after sham or TAC surgery. (D) Representative images of 12-month-old hearts from both genotypes and both surgeries at the 8-week post-surgery collection timepoint. (E) Heart weight to body weight ratios of 6-month-old and (F) 12-month-old animals after TAC or sham surgery. P values indicated above bar graphs are calculated from unpaired t tests. 2-way ANOVA tests were performed on the same cohort of animals before and after their respective surgeries to determine impact of the genotypes on the resulting echocardiography results.

To assess ventricular structure and function, we next compared fractional shortening (FS), left ventricle dimensions in systole and diastole, and the mitral annular tissue velocity (E/E’), which is the diastolic left ventricle filling pressure ratio. In 6-month-old animals, TAC surgery resulted in decreased FS and increased E/E’ in WT hearts, indicative of impaired contractility. *Ficd^-/-^*animals did not show significant changes in FS or E/E’ after TAC surgery compared to their pre-TAC cardiac function (Supplemental Fig. S1C). Because increased age is a leading risk factor of cardiac stress and disease, we also assessed the effects of TAC and sham surgeries on 12-month-old mice. TAC surgery induced a significantly quicker decline in EF in the 12-month-old WT animals compared to the 6-month-old WT animals (P_int_ = 0.009) (Supplemental Fig. S1E). In 12-month-old mice, FICD deficiency was again sufficient to prevent TAC-induced declines in EF at all time points (P_int_ = 0.007) (Fig. 1C, Supplemental Fig. S1A). Consistently, 12-month-old *Ficd^-/-^* animals were protected from a decline in FS after TAC treatment, while 12-month-old WT animals experienced a significant decline in FS function after TAC surgery (Supplemental Fig. S1D).

Moreover, we found that TAC surgery led to an increase in heart mass relative to body mass (HW/BW) in 8-month-old WT hearts (Fig. 1E). Cardiac hypertrophy was significantly reduced in these *Ficd^-/-^* mice after TAC compared to WT animals who underwent TAC surgery (P_int_ = 0.014). Similar to the younger cohort, 14-month-old WT hearts showed an increased HW/BW ratio compared to the HW/BW of *Ficd^-/-^* mice after TAC surgery (Fig. 1D,F, Supplemental Fig. S2). These data suggest that FICD deficiency is protective against pressure-induced hypertrophic stress and preserves cardiac function.

### FICD deficiency reduces cardiomyocyte hypertrophy and fibrosis following pressure overload stress

Pressure overload leads to cardiomyocyte hypertrophy and fibrotic tissue accumulation, resulting in heart enlargement. However, these early compensatory measures cause heart muscle rigidity and muscle loss over time, ultimately leading to heart failure ^2^. We quantified fibrotic tissue accumulation and the size of cardiomyocytes from cryosections of sham and TAC hearts from both genotypes. We observed a significant increase in fibrotic tissue staining in WT hearts that had undergone TAC surgery in both the 6-month-old and 12-month-old experimental cohorts (Fig. 2A-C, Supplemental Fig.S3A). However, cardiac tissue fibrosis was significantly suppressed in the *Ficd^-/-^* hearts from the 12-month-old mice that underwent TAC-induced pressure overload (P_int_ = 0.03) (Fig. 2A-C). Hearts of 6-month-old *Ficd^-/-^* mice post TAC surgery also showed a slight reduction in fibrosis (P_int_ = 0.1). We further observed that TAC surgery increased cardiomyocyte diameter in both genotypes; however this effect was significantly reduced in *Ficd^-/-^*hearts (Fig. 2D,E, Supplemental Fig.S3B). These results further corroborate that FICD deficiency significantly reduces cardiomyocyte hypertrophy upon pressure overload, while not completely abolishing it. Taken as a whole, our results define a novel link between FICD, cardiomyocyte hypertrophy, and cardiac fibrosis in the context of cardiac adaptation to pressure overload.

**Figure 2.**
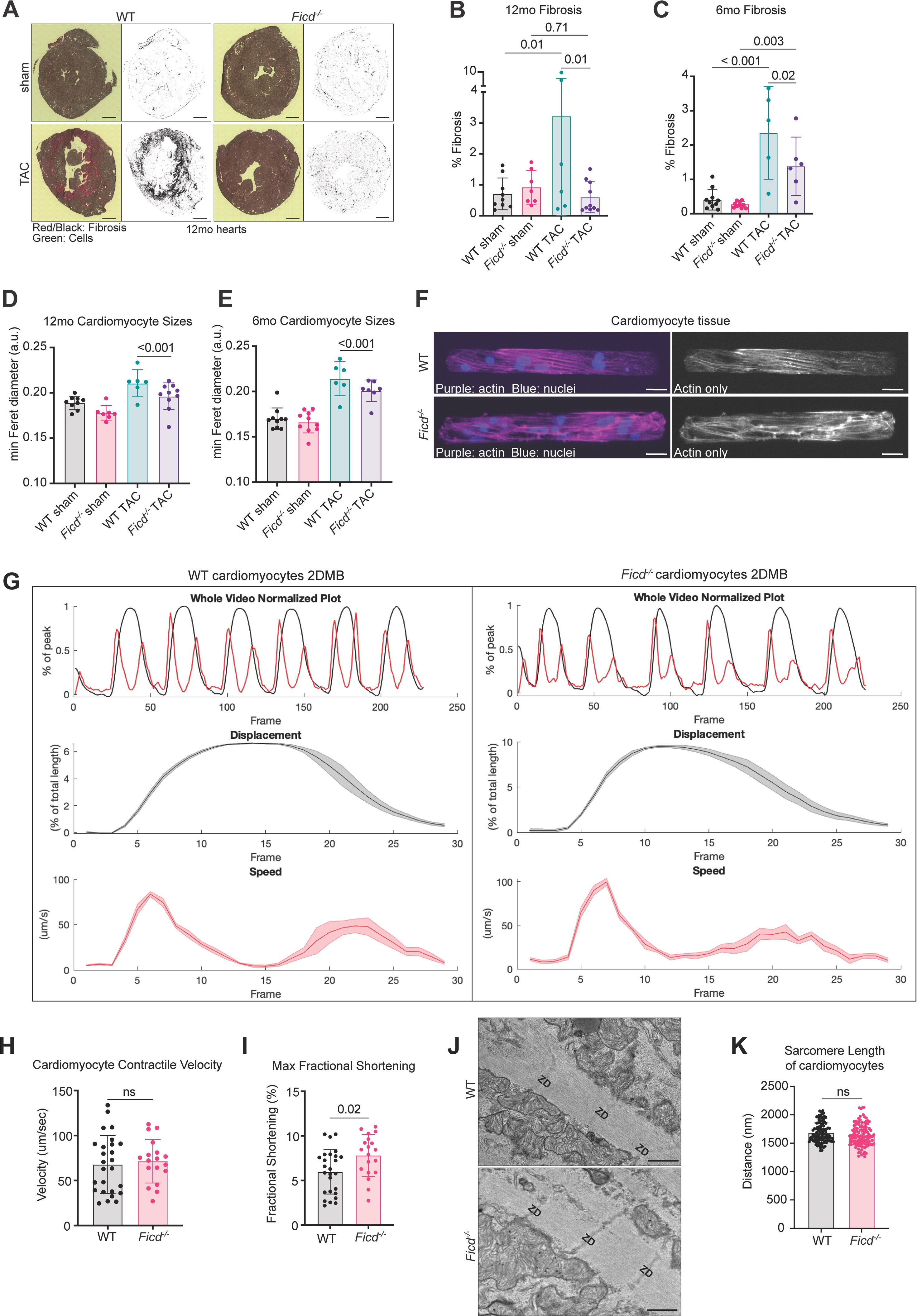
Cardiac Cellular Analysis of WT and *Ficd^-/-^* animals. (A) Representative images of fibrosis (red/black) and cells (green) from cryosectioned hearts after undergoing sham or TAC surgery (Scale bar, 2mm). (B) Quantification of the percent fibrotic tissue detected from cryosections of 12-month mouse (n= 7-12) hearts and (C) 6-month mouse (n= 8-10 per genotype per surgery) hearts after undergoing sham or TAC surgery. (D) Quantification of cardiomyocyte hypertrophy using Feret’s minimum diameter of individual cardiomyocytes from cryosections of 12-month mouse (n= 7-12) hearts and (E) 6-month-old mouse (n= 8-10 per genotype per surgery) hearts after undergoing sham or TAC surgery. P values above bars are from unpaired t tests comparing Feret’s minimum diameter of the individual cardiomyocytes from the cryosections. (F) Isolated primary cardiomyocytes were plated onto 2DMB plates and self-organized organized into ∼7:1 aspect ratio cardiac muscle bundles, each comprised of 10-12 aligned cardiomyocytes. Cells were fixed and stained with alpha-actinin (pink/black and white) and DAPI (blue) to visualize sarcomere and nuclear structures. (Scale bar, 40μm). (G) Representative data from ContractQuant method from MATLAB for fractional shortening (2^nd^ row), velocity measurements (third row), and overlay of both during the (250 frames/5 seconds) acquisition time frame (top row). The solid lines in the graphs are the mean of the values and the shaded area is the 95% confidence interval. (H) Quantification of contractile velocity (μm/sec) and (I) fractional shortening (%) of spontaneously contracting WT and *Ficd^-/-^* cardiac muscle bundles from live cell imaging. (J) Transmission Electron Microscopy (TEM) images of sarcomeres from isolated WT and *Ficd^-/-^* cardiomyocytes. ZD: Z-Disc (Scale bar, 800nm). (K) Quantification of the distance between Z lines in muscle fibers, representing the length of individual sarcomeres, from TEM images of WT and *Ficd^-/-^* isolated cardiomyocytes. Indicated P values were calculated using unpaired t tests. 2-way ANOVA tests were performed in parallel to confirm significance of results.

We next tested whether FICD deficiency acutely affects cardiomyocyte contractility and structure prior to age- or TAC-related remodeling. To answer this question, we used a micron-scale 2D cardiac muscle bundle (2DMB) approach, which enables isolated neonatal cardiomyocytes to arrange in individual muscle strips along micropatterned 7:1 eccentric rectangles. This approach facilitates physiologic alignment and uniaxial contractile direction of immature cardiomyocytes to improve reproducibility of contractility quantification (Supplemental Fig.S4). Primary WT and *Ficd^-/-^* neonatal cardiomyocytes were plated onto the 2DMB platform, where approximately 10-12 cardiomyocytes self-organized into spontaneously contracting 2DMBs with quantifiable contractile rates and distances. Contractile velocity was unchanged in *Ficd^-/-^* 2DMBs (Fig. 2G,H). However, when assessing the displacement of the 2DMBs during contractions, the *Ficd^-/-^*2DMBs had increased fractional shortening (Fig. 2G,I). These data, along with the increased fractional shortening observed in the *Ficd^-/-^*hearts by echocardiography (Supplemental Fig.1C,D), suggest a potential link between FICD activity and cardiomyocyte contractility even in the absence of pressure overload. Using immunofluorescence and transmission electron microscopy (TEM), we further observed that structural components, including contractile machinery and membrane of WT and *Ficd^-/-^* cardiomyocytes and cardiac fibroblasts remained consistent (Fig. 2F,J,K, Supplemental Fig. S4). These results suggest that FICD loss improves contractility in 2DMBs even in the absence of hypertrophic stimuli.

### The impact of ER stress on cardiac FICD activity is cell-type specific

BiP is an ER-resident HSP70 chaperone regulated by FICD-mediated AMPylation. Since BiP acts as a major regulator of the UPR^ER^, we sought to understand how FICD activity contributes to ER stress resilience in cardiac tissue. We first assessed BiP AMPylation in hearts of animals after TAC or sham surgery. As expected, FICD deficient hearts did not show any BiP AMPylation signature under any experimental condition (Fig. 3A-D,G-H, Supplemental Fig. S5). In contrast, WT hearts collected 2 months post-TAC surgery presented with increased BiP AMPylation compared to the WT sham controls (Fig. 3E-F, Supplemental Fig. S5). These results suggest that chronic pressure overload increases BiP AMPylation and has lasting impacts on ER proteostasis regulation. To understand how individual cardiac cell-types react to ER stress, we isolated primary mouse cardiomyocytes and cardiac fibroblasts and treated both cell types with the glycosylation inhibitor tunicamycin (Tm) or the Sarcoplasmic/Endoplasmic Reticulum Ca2+-ATPase (SERCA) inhibitor thapsigargin (Tg), both of which induce ER stress. The chemical induction of ER stress increased BiP protein levels in cultured cardiomyocytes (Fig. 3I-L). We also observed ER stress significantly reduced BiP AMPylation in cardiomyocytes (Fig. 3I-J). However, when isolated cardiac fibroblasts were exposed to the same two stressors, only Tg treatment led to a significant decrease in BiP AMPylation; whereas Tm treatment significantly increased BiP AMPylation relative to untreated cardiac fibroblasts (Fig. 3K-L). These results indicate that ER stress-induced FICD activity is cell type-specific.

**Figure 3.**
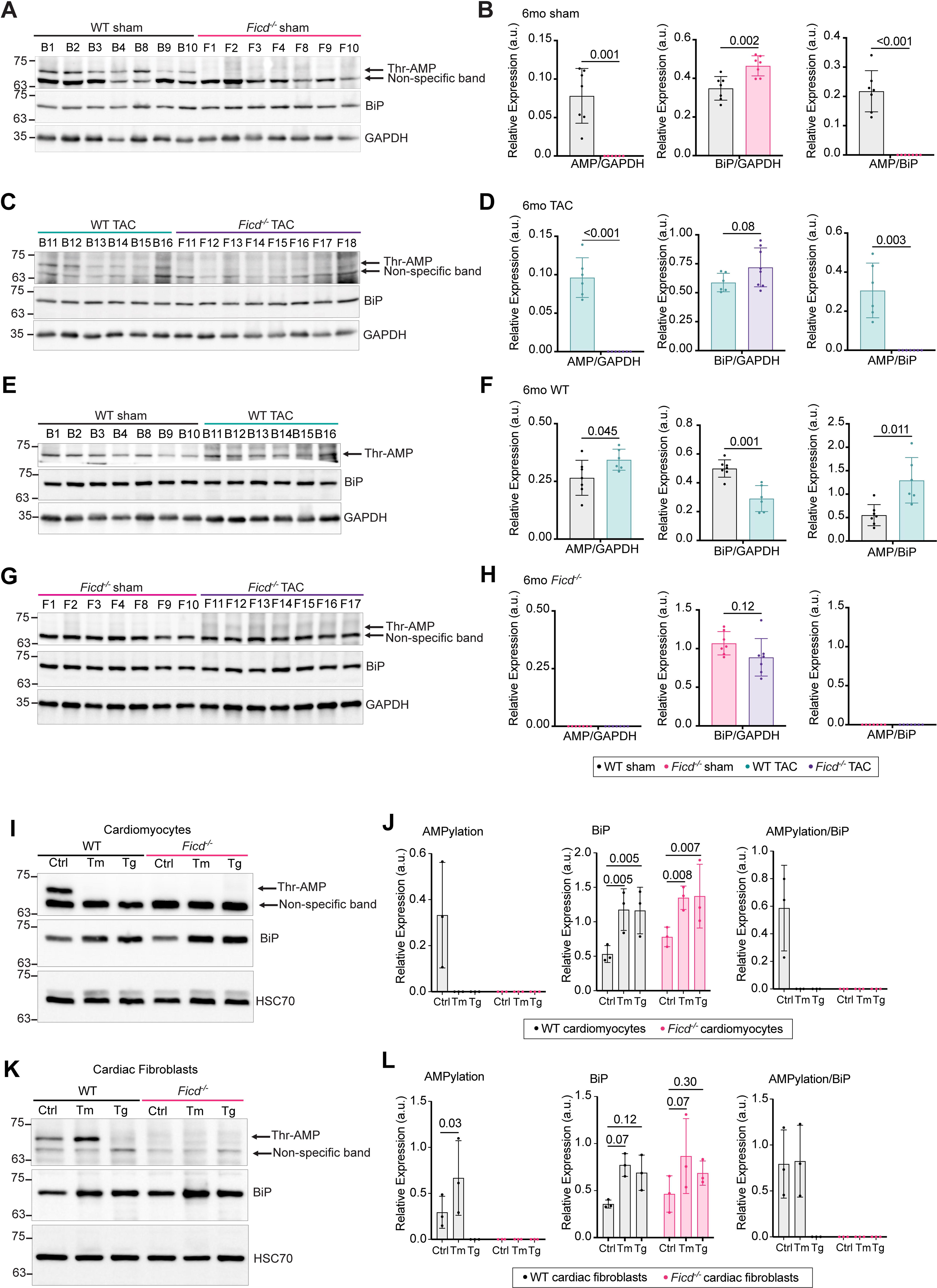
BiP AMPylation is differentially regulated by FICD in Cardiomyocytes and Cardiac Fibroblasts. Hearts were collected from 8-month mice 8 weeks post-TAC or sham surgery and processed for protein quantification. Protein from (A) sham only, (C) TAC only, (E) WT only, and (G) *Ficd^-/-^*only mice were run on 10% Tris-Glycine SDS acrylamide gels, blocked with 5% BSA, and probed for Thr-AMPylation, BiP, and GAPDH (loading control) levels (n=6-7 mice per genotype per condition). Quantification of Thr-AMPylation, BiP, and GAPDH from (B) sham only, (D) TAC only, (F) WT only, (H) *Ficd^-/-^*only blots. (I) Primary cardiomyocytes and (K) primary cardiac fibroblasts were isolated from 6-8 mice hearts per condition and plated in 6-well dishes for 48 hours at 37°C and 5% CO_2_. Cardiomyocytes and cardiac fibroblasts were treated with Tm (10μg/mL) or Tg (3μM) for 12 hours before being collected for protein quantification via western blotting. Cells were run on 10% Tris-Glycine SDS acrylamide gels, blocked with 5% BSA, and probed for Thr-AMPylation, BiP, and HSC70 (loading control) levels. (J) Levels of Thr-AMPylation, BiP, and HSC70 were quantified from control and ER-stressed cardiomyocytes and (L) cardiac fibroblasts. Indicated P values were calculated using unpaired t tests. 2-way ANOVA tests were performed in parallel to confirm significance of results.

### FICD mitigates the UPR^ER^ in stressed cardiac tissues

Because the response of BiP AMPylation to ER stress was cell-type specific, we hypothesized that FICD loss would have distinct consequences on UPR^ER^ signaling in cardiomyocytes and cardiac fibroblasts. Indeed, we found that isolated *Ficd^-/-^* cardiomyocytes showed a more robust activation of all three arms of the UPR^ER^ when stressed with Tm or Tg relative to WT cardiomyocytes under the same stress (Fig. 4A-D). *Ficd^-/-^* cardiac fibroblasts treated with Tm or Tg showed a significant increase in the ratio of spliced versus non-spliced *Xbp-1*, indicating enhanced activation of the IRE-1 arm of the UPR^ER^ (Fig. 4E). In contrast, PERK-dependent upregulation of *Atf-4* in response to ER stress was not elevated in FICD deficient cardiac fibroblasts compared to controls (Fig. 4F). Atf6α transcription increased in fibroblasts of both genetic backgrounds in the presence of ER stressors, albeit significantly more in Tg-treated *Ficd^-/-^* cells (Fig. 4G). Next, we assessed how FICD loss alters expression of C/EBP Homologous Protein (CHOP), a mediator of ER-induced apoptosis. We observed that Tm and Tg treatment both increased CHOP expression in WT cardiomyocytes. However, *Ficd^-/-^*cardiomyocytes showed a significant reduction in CHOP expression in response to Tg compared to WT controls (Fig. 4I-J). Contrastingly, cardiac fibroblasts displayed a robust increase in CHOP expression in response to either ER stressor in both WT and *Ficd^-/-^*backgrounds (Fig. 4K-L). These results suggest that in the absence of FICD, cardiomyocytes increase activity of all branches of the UPR^ER^ during ER stress. Meanwhile, in cardiac fibroblasts, the UPR^ER^ response is restricted to IRE1α and Tg-induced ATF6 activation.

**Figure 4.**
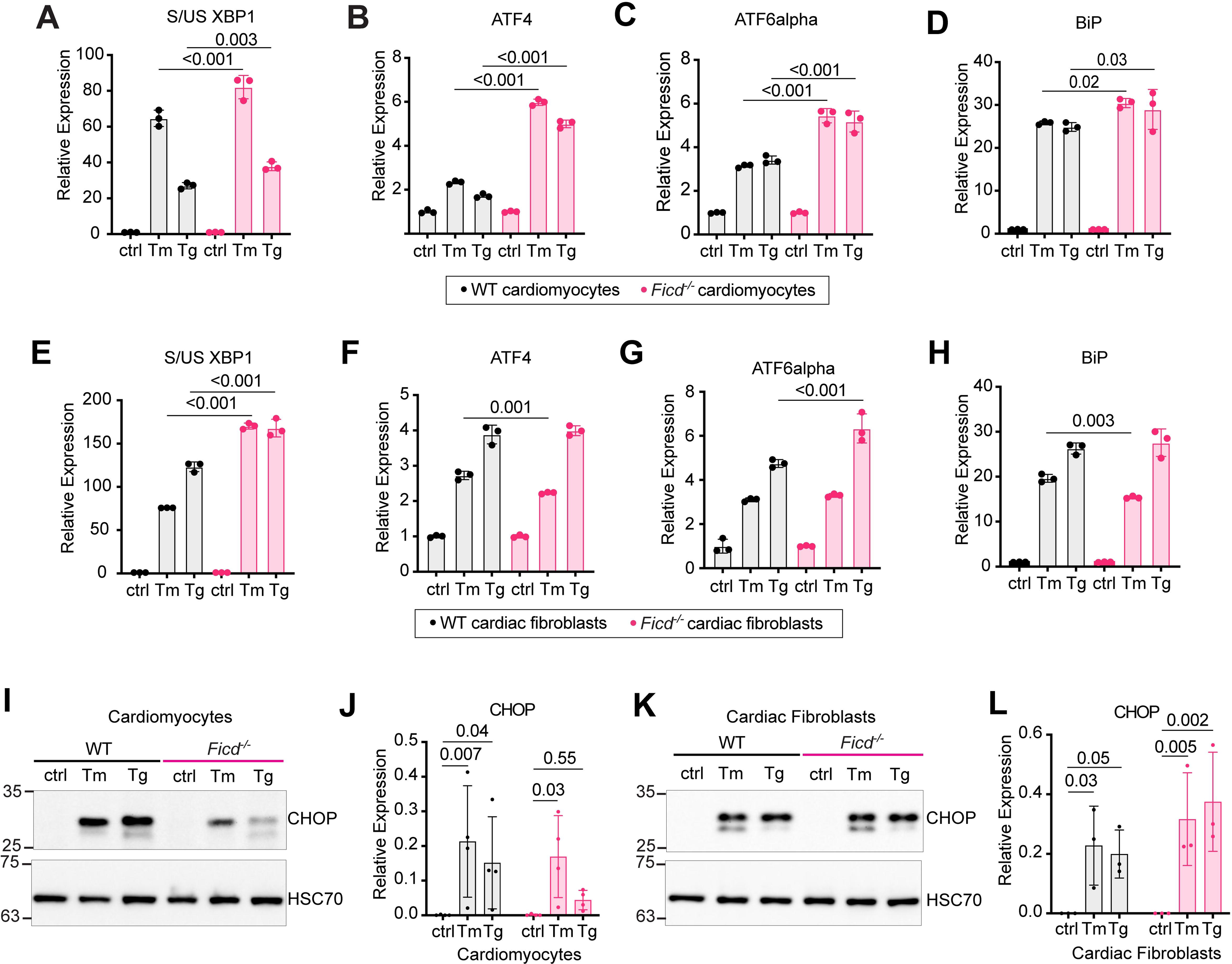
Different branches of the UPR^ER^ are activated in *Ficd^-/-^* cardiomyocytes and cardiac fibroblasts under ER stress. Real time quantitative PCR on mRNA from isolated WT and *Ficd^-/-^*cardiomyocytes treated with control, Tm (10μg/mL), or Tg (3μM) media, assessed for amounts of (A) S/US XBP1, (B) ATF4, (C) ATF6alpha, and (D) BiP. Real time quantitative PCR on mRNA from isolated WT and *Ficd^-/-^* cardiac fibroblasts treated with control, Tm (10μg/mL), or Tg (3μM) media, assessed for amounts of (E) S/US XBP1, (F) ATF4, (G) ATF6alpha, and (H) BiP. Gene expression is normalized to the respective control group of each genotype. Protein from (I) cardiomyocytes and (K) cardiac fibroblasts were run on 10% Tris-Glycine SDS acrylamide gels, blocked with 5% BSA, and probed for CHOP and HSC70 (loading control) levels. Levels of CHOP and HSC70 were quantified from control and ER-stressed (J) cardiomyocytes and (L) cardiac fibroblasts. Indicated P values were calculated using unpaired t tests. n= 6-8 mice per condition.

### FICD limits ER-stress induced autophagy

Pressure-induced hypertrophy requires cardiac tissues to increase protein recycling through the autophagosome-lysosomal pathway^48,85,86^. Because ER stress can induce autophagy through the UPR^ER^, we predicted that FICD deficiency would lead to enhanced autophagic protein clearance. We thus evaluated the abundance of lipidated LC3-II, representative of autophagosome abundance, in 6-month-old hearts collected post sham or TAC surgery. We found that WT and *Ficd^-/-^* sham hearts did not exhibit differences in LC3-II levels (Fig. 5A,E). In contrast, TAC-induced pressure overload led to a significant increase in LC3-II levels in *Ficd^-/-^* hearts relative to WT controls (Fig. 5B,F). Consistent with previous studies demonstrating impaired autophagy in hypertrophic hearts^87^, we also found that LC3-II levels decreased in WT hearts following pressure overload, yet LC3-II levels remained unchanged in *Ficd^-/-^* hearts after TAC (Fig. 5C,D,G,H). These results demonstrate that the presence of autophagosomes is unchanged before and after hypertrophic stress in the absence of FICD, while WT hearts experience a reduction of these vesicles following hypertrophic stress.

**Figure 5.**
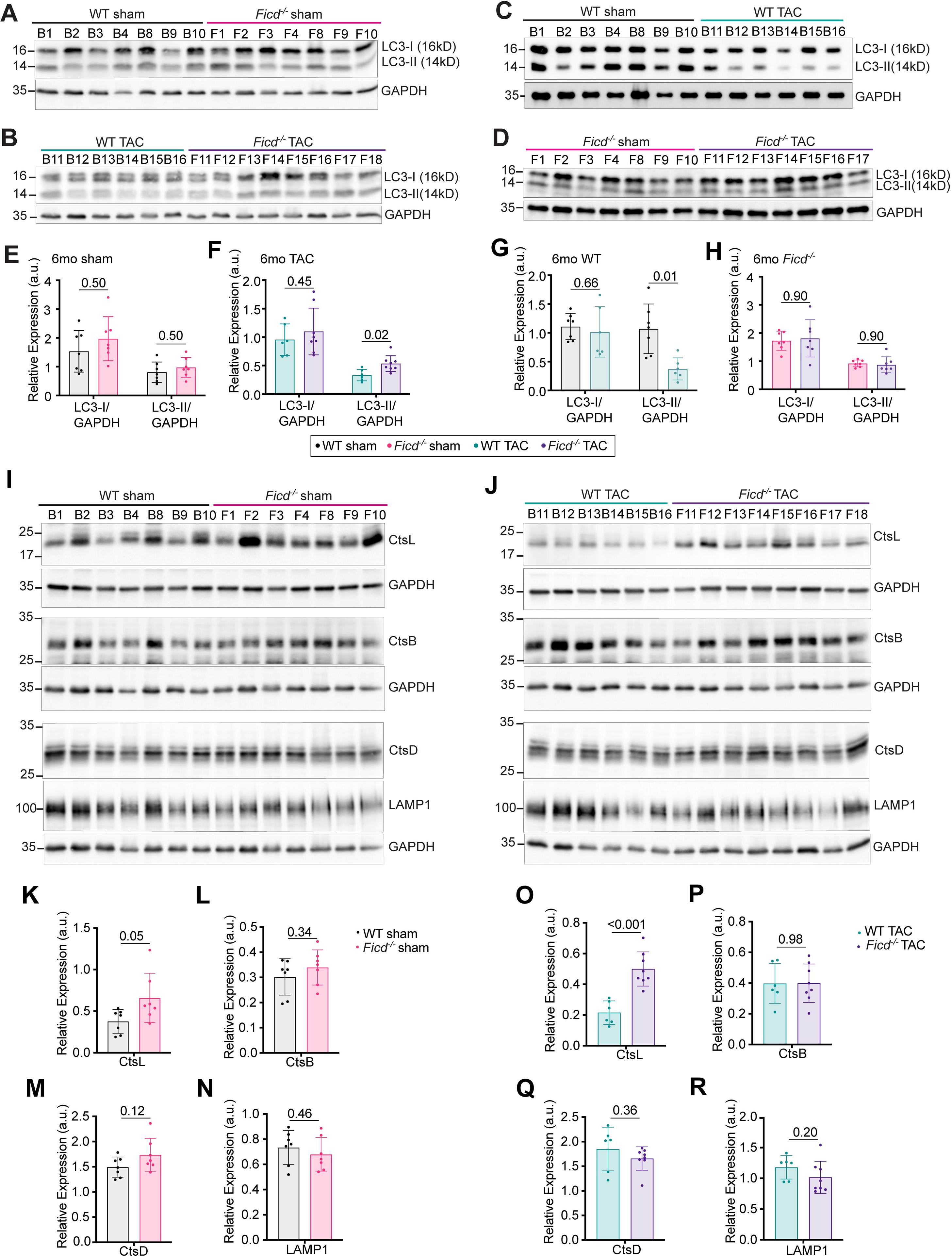
LC3-II and Cathepsin L are upregulated after TAC surgery in *Ficd^-/-^* hearts. Hearts were collected from 8-month mice 8 weeks post-TAC or sham surgery and processed for protein quantification. Protein from (A) sham only, (B) TAC only, (C) WT only, and (D) *Ficd^-/-^* only mice were run on 15% Tris-Glycine SDS acrylamide gels, blocked with 5% BSA, and probed for LC3 and GAPDH (loading control) levels (n=6-7 mice per genotype per condition). Quantification of LC3-I, LC3-II, and GAPDH from (E) sham only, (F) TAC only, (G) WT only, (H) *Ficd^-/-^*only gels. Protein from (I) sham only and (J) TAC only mice were run on 10% Tris-Glycine SDS acrylamide gels, blocked with 5% BSA, and probed for CtsL, CtsB, CtsD, LAMP1, and GAPDH (loading control) levels (n=6-7 mice per genotype per condition). Quantification of (K) CtsL, (L), CtsB, (M) CtsD, and (N) LAMP1 levels relative to GAPDH controls in sham only mice. Quantification of (O) CtsL, (P), CtsB, (Q) CtsD, and (R) LAMP1 levels relative to GAPDH controls in TAC only mice. Indicated P values were calculated using unpaired t tests.

Increased LC3-II levels can be attributed to an increase in autophagy induction or a decrease in lysosomal processing of autophagosomes. To determine if FICD deficiency alters lysosomal activity, we evaluated levels of lysosomal proteins in WT and *Ficd^-/-^*hearts collected post-sham or TAC surgery. We assessed protein levels of the cathepsin family, a group of lysosomal proteases that function to degrade protein cargo delivered from autophagosomes to lysosomes^88–91^. We found that FICD loss significantly increased basal Cathepsin L (CtsL) expression but did not affect levels of Cathepsins B or D (CtsB and CtsD) (Fig. 5I, K-N). Pressure overload further enhanced differences in CtsL expression between FICD deficient and control hearts while CtsD, CtsB, and LAMP1 expression remained constant (Fig. 5J,O). These results demonstrate that lysosomal activity in FICD deficient hearts is likely not impaired due to consistent changes in lysosomal protein abundance. Further, they support the conclusion that the observed increase in LC3-II levels in *Ficd^-/-^*TAC hearts results from an increase in autophagy activity rather than a decrease in lysosomal processing. Our results further provide the first evidence for a direct link between FICD and autophagy while suggesting that *Ficd^-/-^* animals may be protected from hypertrophic-induced heart failure by the activation of the autophagy-lysosomal pathway.

### FICD-mediated BiP AMPylation prevents BiP-FAM134B interactions thus blocking ER-phagy

To confirm that FICD deficiency improves autophagic processing, we performed autophagy flux assays in isolated cardiomyocytes in the presence of ER stress. We treated WT and *Ficd^-/-^* cardiomyocytes with combinations of Tm, Tg, and bafilomycin, an inhibitor of autophagosome-lysosome fusion. Bafilomycin treatment significantly increased LC3-II levels in WT and *Ficd^-/-^* cardiomyocytes, indicative of active autophagic processes. This autophagic flux was maintained in both genotypes in the presence of ER stressors (Fig. 6A-B). Since autophagic flux depends on lysosomal function, we evaluated cathepsin levels in isolated cardiomyocytes exposed to ER stress. Consistent with our *in vivo* heart data, we found that CtsL was significantly upregulated in *Ficd^-/-^* cardiomyocytes both under basal and ER stress conditions (Fig. 6E,F). Protease activity assays further confirmed that *Ficd^-/-^* cardiomyocytes were equipped with significantly enhanced CtsL activity in the absence of ER stress (Fig. 6G). In contrast, CtsB, CtsD, and LAMP1 expression remained unaffected between WT and *Ficd^-/-^* cells in the absence of ER stress. The induction of ER stress using Tg, but not Tm, resulted in a significant boost in CtsB, CtsD, and LAMP1 abundance in *Ficd^-/-^*cardiomyocytes (Fig. 6E, H-J). Taken as a whole, these results confirm that FICD deficiency increases autophagic flux in cardiomyocytes, possibly through an increase in CtsL abundance and activity.

**Figure 6.**
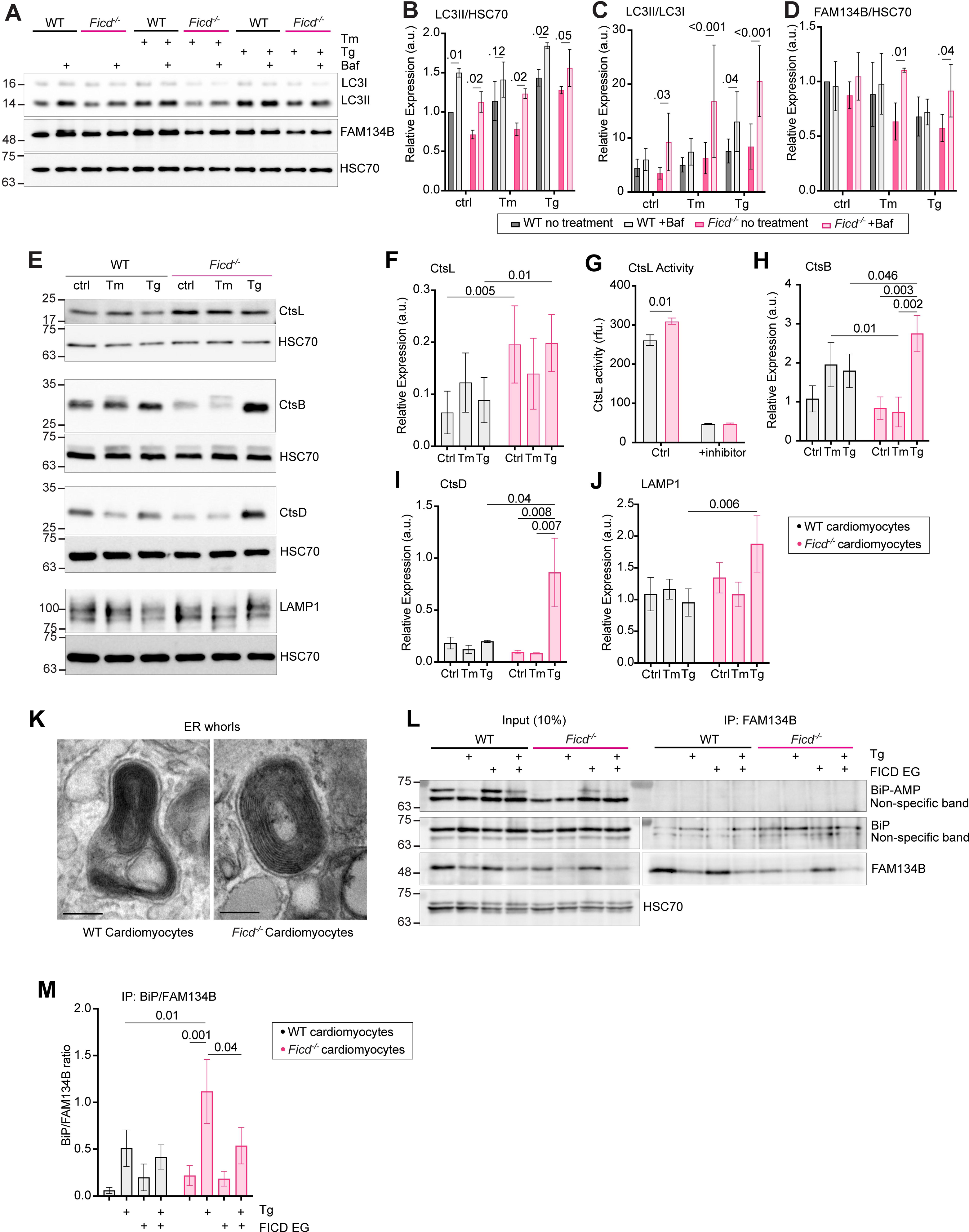
ER-phagy is activated via a BiP/FAM134B interaction in *Ficd^-/-^* cardiomyocytes under stress. (A) Primary WT and *Ficd^-/-^* cardiomyocytes were isolated from 6-8 mice hearts per condition and plated in 24-well dishes for 48 hours at 37°C and 5% CO2. Cardiomyocytes were treated with Tm (10μg/mL) or Tg (3μM) for 10 hours and supplemented with bafilomycin (100nM) for four hours before being collected for protein quantification via western blotting. Cells were run on 15% Tris-Glycine SDS acrylamide gels, blocked with 5% BSA, and probed for LC3, FAM134B, and HSC70 (loading control) levels. Levels of (B) LC3-II/HSC70, (C) LC3-II/LC3-I, and (D) FAM134B/HSC70 were quantified from control, ER-stressed, and bafilomycin treated cardiomyocytes. (E) Primary WT and *Ficd^-/-^* cardiomyocytes were isolated from 6-8 mice hearts per condition and plated in 6-well dishes for 48 hours at 37°C 5% CO2. Cardiomyocytes were treated with Tm (10μg/mL) or Tg (3μM) for 12 hours before being collected for protein quantification via western blotting. Cells were run on 10% Tris-Glycine SDS acrylamide gels, blocked with 5% BSA, and probed for CtsL, CtsB, CtsD, LAMP1, and HSC70 (loading control) levels. Levels of (F) CtsL/HSC70, (H) CtsB/LC3-I, (I) CtsD/HSC70, and (J) LAMP1/HSC70 were quantified from control and ER-stressed cardiomyocytes. (G) CtsL activity from 50μg of isolated WT and *Ficd^-/-^* cardiomyocytes from non-treated and CtsL inhibitor treated cells. (K) Transmission Electron Microscopy images of ER whorls from isolated WT and *Ficd^-/-^* cardiomyocytes. (Scale bar, 200nm). (L) Co-immunoprecipitation of FAM134B-BiP from control, Tg-treated, and FICD(E324G) transfected WT and *Ficd^-/-^* cardiomyocytes. FAM134B antibody was incubated with 150μg of protein from each cardiomyocyte condition and immunoprecipitated with Protein A Magnetic agarose beads. 10% input and the IP-ed proteins were run on 10% Tris-Glycine acrylamide gels, blocked with 5% BSA, and probed for Thr-AMPylation, BiP, FAM134B, and HSC70 (loading control). (M) Quantification of three replicate co-immunoprecipitations of BiP/FAM134B from control, Tg, and FICD(E324G) transfected WT and *Ficd^-/-^* cardiomyocytes. Indicated P values were calculated using unpaired t tests. 2-way ANOVA tests were performed in parallel to confirm significance of results.

We found that lysosome inhibition significantly increased the LC3-II/LC3-I ratio in *Ficd^-/-^* cardiomyocytes, particularly in the presence of ER stress (Fig. 6C). While not indicative of total autophagosome abundance, these data could signify an increase in autophagosome formation in the absence of FICD. Further, when assessing cardiomyocytes using TEM, we observed whorl structures in both WT and *Ficd^-/-^* cardiomyocytes (Fig. 6K). Whorls typically form during ER-phagy due to engulfed ER membranes stacking closely together in the autophagosome^92^. ER-phagy plays an important role in degrading cytotoxic portions of the ER to recover cellular homeostasis. To discern if FICD was regulating autophagy initiation through ER-phagy, we assessed the abundance of FAM134B, a well-known ER-phagy receptor, in WT and *Ficd^-/-^* cardiomyocytes. FAM134B is lysosomally degraded during ER-phagy. Therefore, we measured ER-phagy activity by quantifying FAM134B levels during the previously described flux assay. We found that FAM134B significantly accumulated in the presence of ER stress and bafilomycin in *Ficd^-/-^* cardiomyocytes (Fig. 6D). This suggests that ER stress leads to ER-phagy-mediated degradation of FAM134B in *Ficd^-/-^* cardiomyocytes, in contrast, FAM134B flux remained unchanged in WT cardiomyocytes.

BiP initiates ER-phagy upon binding to the ER luminal domain of FAM134B ^55^. We thus hypothesized that a lack of FICD-mediated BiP AMPylation could lead to enhanced BiP engagement of FAM134B in FICD deficient cardiomyocytes. To test this, we assessed the impact of BiP AMPylation on its ability to bind to FAM134B. Since ER stress leads to BiP deAMPylation (Fig. 3A), we over-expressed a constitutive-active AMPylase mutant (FICD E234G) in cardiomyocytes, allowing us to maintain a fraction of AMPylated BiP under ER stress (Fig. 6L). We immunoprecipitated FAM134B from these cardiomyocytes and confirmed that BiP co-immunoprecipitated with FAM134B (Fig. 6L,M). Notably, only non-AMPylated BiP interacted with FAM134B under ER-phagy-initiating conditions (Fig. 6L,M). Further, we observed an increase in FAM134B-mediated BiP pulldown in *Ficd^-/-^*cardiomyocytes under ER stress, but this effect was suppressed in the presence of FICD(E234G) (Fig. 6L,M). These results support the model that FICD-mediated BiP AMPylation acts as a gatekeeper for ER-phagy induction in the presence of ER stress in cardiomyocytes, and that only upon the deAMPylation of BiP can the BiP/FAM134B complex form to initiate ER-phagy.

## Discussion

In this study, we demonstrate that FICD deficiency prevents a decline in cardiac ejection fraction following TAC-induced pressure overload. We further show that FICD loss mitigates *in vivo* hypertrophic growth and fibrotic tissue accumulation in the heart. Mechanistically, we link cell type-specific FICD activity to the regulation of the UPR^ER^ and ER-phagy through BiP AMPylation and FAM134B engagement (Fig. 7). In the absence of FICD, BiP is not AMPylated in cardiomyocytes to alleviate ER stress via increased activation of UPR^ER^ and ER-phagy.

**Figure 7.**
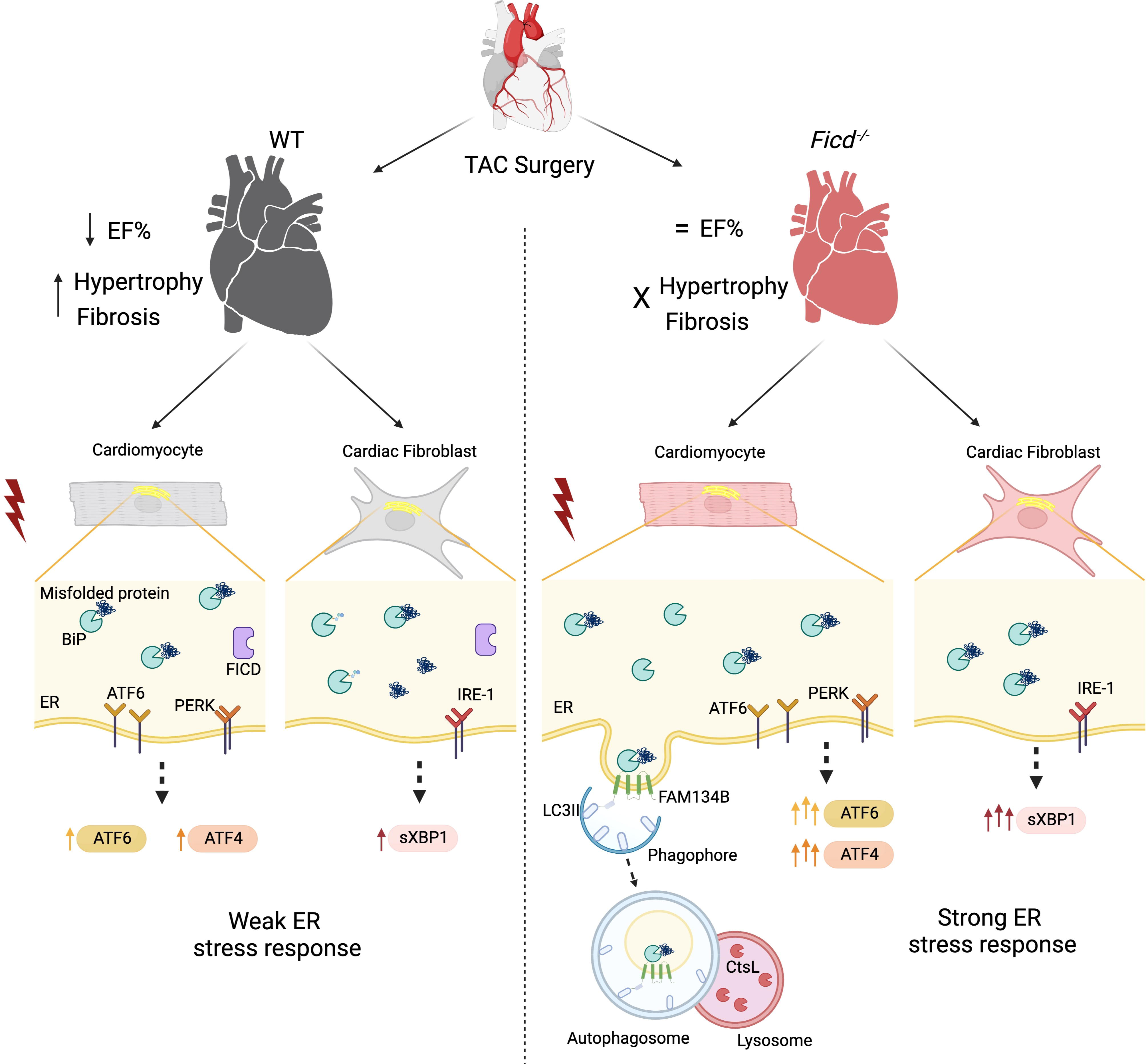
Model of molecular mechanisms underlying the protective phenotype of *Ficd^-/-^*animals under TAC surgery. Representation of the different ER stress responses between WT and *Ficd^-/-^* hearts to pressure-induced hypertrophy in cardiomyocytes and cardiac fibroblasts. Grey cells reflect cells with weaker proteostasis response and pink cells reflect cells with stronger proteostasis response. ER stress-induced misfolded proteins in *Ficd^-/-^* cells are efficiently ameliorated by the BiP/FAM134B initiated ER-phagy response and enhanced upregulation of the UPR^ER^ compared to WT cells.

Assessing cardiomyocytes and cardiac fibroblasts, two predominant cardiac-resident cell types, we show that FICD-deficient cardiac cells exhibit a more robust induction of the UPR^ER^ in response to ER stress. UPR^ER^ induction is required to support recovery from acute ER stress, but prolonged UPR^ER^ signaling can lead to ER-mediated apoptosis. Previous studies using mouse *Ficd^-/-^* embryonic fibroblasts, FICD-ablated AR42 cells, and HEK293T cells treated with *Ficd* siRNA highlight direct links between FICD activity and the induction of the UPR^ER^^58,68,93^. In all examples, ER stress in *Ficd* deficient cells led to the upregulation of UPR^ER^ genes, albeit often at lower levels than in control cells. Further, each cell type showed a distinctive pattern of UPR^ER^ induction in the absence of FICD activity. Our work demonstrates that the consequences of FICD deficiency are cell-type specific in the heart and thus may vary even within a single tissue. We show that for the first time, the same ER stress (Tm) has opposite outcomes in cardiomyocytes and cardiac fibroblasts, with Tm inducing deAMPylation of BiP in cardiomyocytes and increasing BiP AMPylation in cardiac fibroblasts. We hypothesize that the contradictory impact on BiP AMPylation may account for the differential UPR^ER^ activity following the same ER stress in different cell types. These results highlight that it is imperative to consider tissue origin and species when studying FICD-mediated UPR^ER^ regulation. While the induction of ER-mediated apoptosis in short-lived cells might be an efficient strategy to maintain functional tissue, long-lived cells, including cardiomyocytes and neurons, depend on mitigating prolonged stress; this observation may explain their ability to tolerate UPR^ER^ induction without inducing apoptosis.

Our experiments show that FICD deficiency leads to enhanced ER-phagy in cardiomyocytes. Consistent with our results, autophagic activity and enhanced CtsL levels have been previously linked to a beneficial cardiac response to hypertrophic stress^94^. Our data further support a model in which BiP AMPylation prevents its interaction with FAM134B, thus inhibiting the induction of ER-phagy through FAM134B activation. BiP-dependent ER-phagy induction is well established: The accumulation of misfolded proteins in the ER promotes the binding of BiP to FAM134B in the ER lumen^55^. This binding allows for the sequestration of proteotoxic entities away from less-affected ER compartments to be degraded via ER-phagy. Our findings support this model, while further suggesting that only non-AMPylated BiP is capable of engaging with FAM134B.

We find for the first time that FICD deficiency increases ER-phagy. Previous studies have primarily focused on FICD-mediated UPR^ER^ regulation^58,68,74^. Our work describes how the AMPylation of BiP controls its ability to bind to FAM134B and thus initiate ER-phagy. We also find Tg-induced increases in CtsL, CtsB, CtsD, and LAMP1 in *Ficd^-/-^* cardiomyocytes. We thus speculate that FICD deficiency increases autophagic protein recycling through at least two mechanisms: enhanced ER-phagy initiation and increased lysosomal activation. Considering that previous studies suggested the presence of AMPylated proteins in the lysosome^95,96^, future studies to understand whether lysosomal protein maturation may be regulated by FICD or if ER-phagy may transfer active FICD to lysosomes are required to better understand this novel signaling paradigm.

Finally, our data shows that in both the intact heart and isolated cultured cardiomyocytes, fractional shortening in FICD deficient cardiomyocytes was increased, suggesting enhanced contractile abilities. This could be due to a change in the myofilament protein turnover resulting in more myofibrils, or to a regulatory mechanism being altered, such as PTMs of myofilament proteins or indirect signaling affecting calcium handling; however this is unknown at this time and requires further study. Previous studies established that BiP is essential to sustain cardiac contractility and health^30^. Our data corroborates these published results and further shows that targeting FICD could represent a novel avenue to mitigate hypertrophy-associated cardiac conditions.

### Limitations of this study

Because we collected hearts eight weeks after pressure overload insult, our results do not capture acute changes in UPR^ER^ and ER-phagy during early cardiac remodeling. Future studies should focus on utilizing cardiomyocyte and cardiac fibroblast restricted *Ficd^-/-^* animal models to demonstrate which cell-specific FICD loss is required to improve cardiac resilience to pressure overload stress. We also cannot exclude that cardiac tissue inflammation may also contribute to the observed improvement in heart function. While our study was not sufficiently powered to assess biological sex as a variable, we did not observe any trends indicating sex-specific differences (Supplemental Fig. S1F).

## Acknowledgments

We thank the members of the Truttmann lab for helpful comments and discussion. William Giblin, Benedict Abdon, and Mary Skinner are acknowledged for proof-reading the manuscript draft. SML was supported by the Cellular and Molecular Biology Program Training Grant T32 GM007315-43/44, as well as by the National Heart, Lung, And Blood Institute of the National Institutes of Health under Award Number F31HL158093. We are responsible for the data presented in this publication and the presented findings are not necessarily representative of the official views of the National Institutes of Health. MCT is supported by an Alzheimer’s foundation Young Investigator Award, a Ruth K Broad foundation award and grant 1R35GM142561. Graphics were created with Biorender.com. We thank the University of Michigan Physiology Phenotyping core members, Steve Whitesall and Kimber Converso-Baran, for performing the sham and TAC surgeries as well as all of the echocardiography assessments. We thank Dorothea Hopfner and Aymelt Itzen for sharing antibody 17G6 to detect AMPylation with us. We thank Joseph Endicott and Jeffrey Knupp for productive discussions. We thank Mary Skinner for teaching us the heart isolation method from neonatal mice and for her input on the published data. We thank Eric Smith and Karen Jin for support using the 2DMB plating method and analysis, as well as supplying us with the 2DMB plates. We thank the University of Michigan Microscopy core, Orthopaedic Research Histology core, Advanced Genomics core, the Miller Lab, and the Pletcher Lab for sharing critical equipment.

## Author contributions

MCT supervised the project. SML, MCT, DM, and ASH planned and designed the experiments. SML, RJT, NDU, SNT, and EDS performed all experiments. SML and MCT wrote the manuscript, and all authors edited and approved the final manuscript.

## Data availability statement

The unprocessed raw datasets (images) generated and analyzed during the current study are available from the corresponding author upon reasonable request.

## Additional information

The authors declare no competing interests.

## Supplementary

**Supplementary Figure S1.**
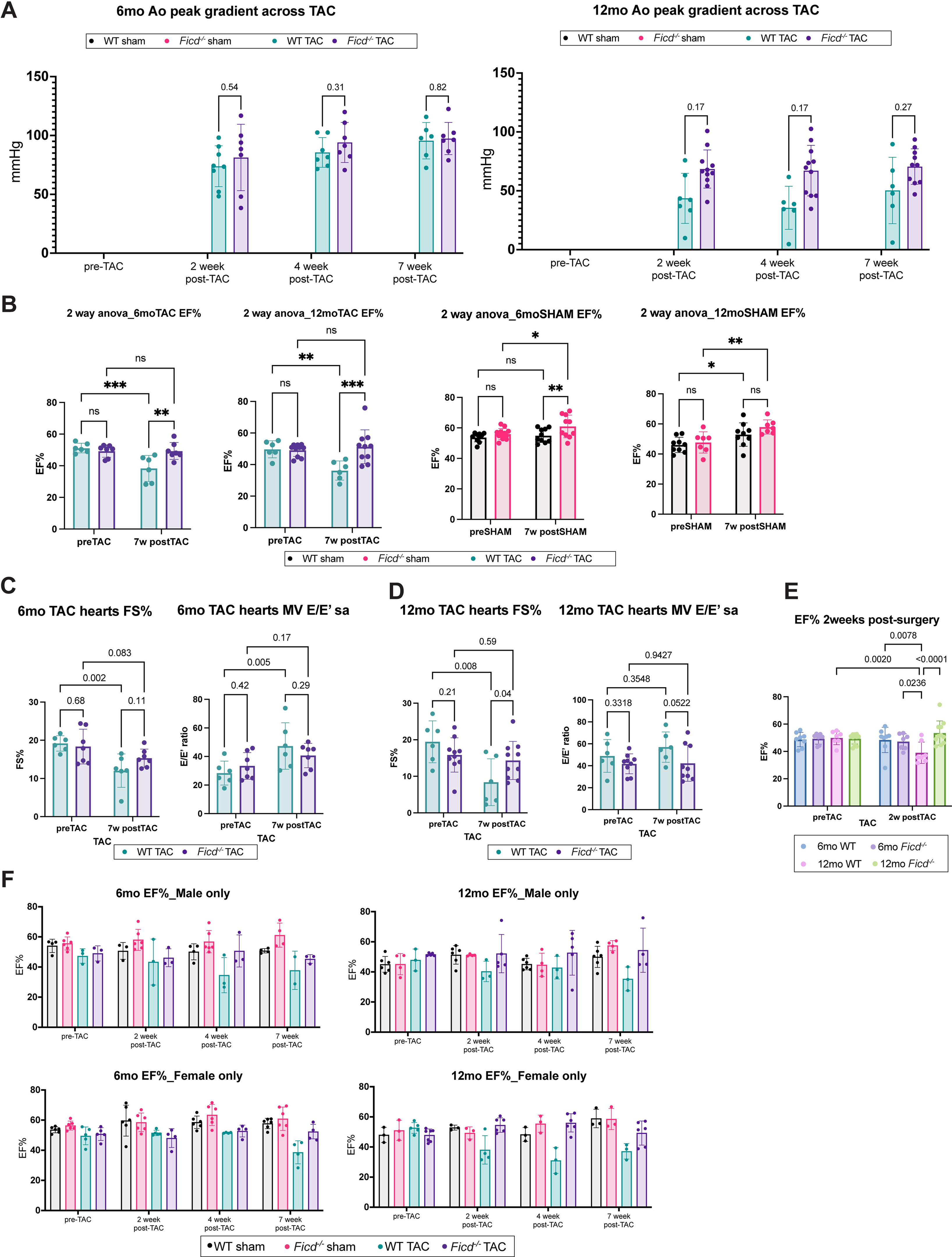
FICD deficient mice are protected from hypertrophic-induced heart failure. (A) Echocardiographic results of the pressure gradients across the TAC ligature on the aortas of 6-month mice (n= 8-10 per genotype per surgery) and 12-month mice (n= 7-12) after sham or TAC surgery. (B) 2-way ANOVA analyses of echocardiographic results of EF pre- and post TAC or sham surgeries in 6-month and 12-month animals. (C) Echocardiographic results of the Fractional Shortening (m-mode short axis imaging) and mitral annular velocity (Doppler imaging) of 6-month mice (n= 8-10 per genotype per surgery) and (D) 12-month mice (n= 7-12) after sham or TAC surgery. (E) Echocardiographic results comparing age, genotype, and surgery impact on EF at the two-week post-TAC timepoint for 3-way ANOVA analysis. P values written above bar graphs are calculated from unpaired t tests. 2-way ANOVA tests were performed on the same cohort of animals before and after their respective surgeries to determine impact of the surgery on resulting echocardiography results. (F) EF echocardiographic results separated by sex of 6-month mice and 12-month mice after sham or TAC surgery.

**Supplementary Figure S2.**
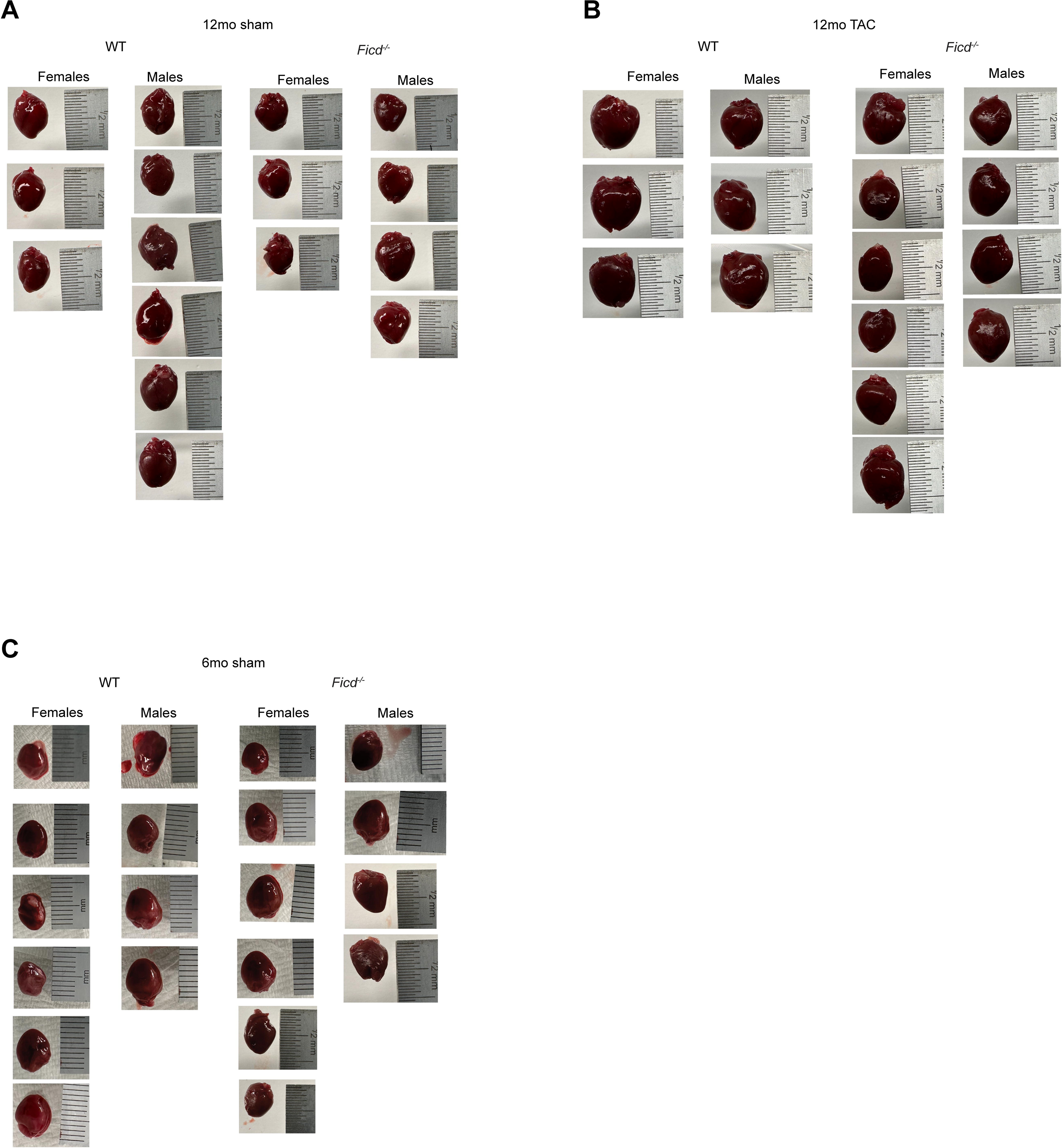
FICD deficient mice are protected from hypertrophic-induced heart failure. Images of collected sham and TAC hearts from 6-month and 12-month animals 8 weeks after respective surgeries.

**Supplementary Figure S3.**
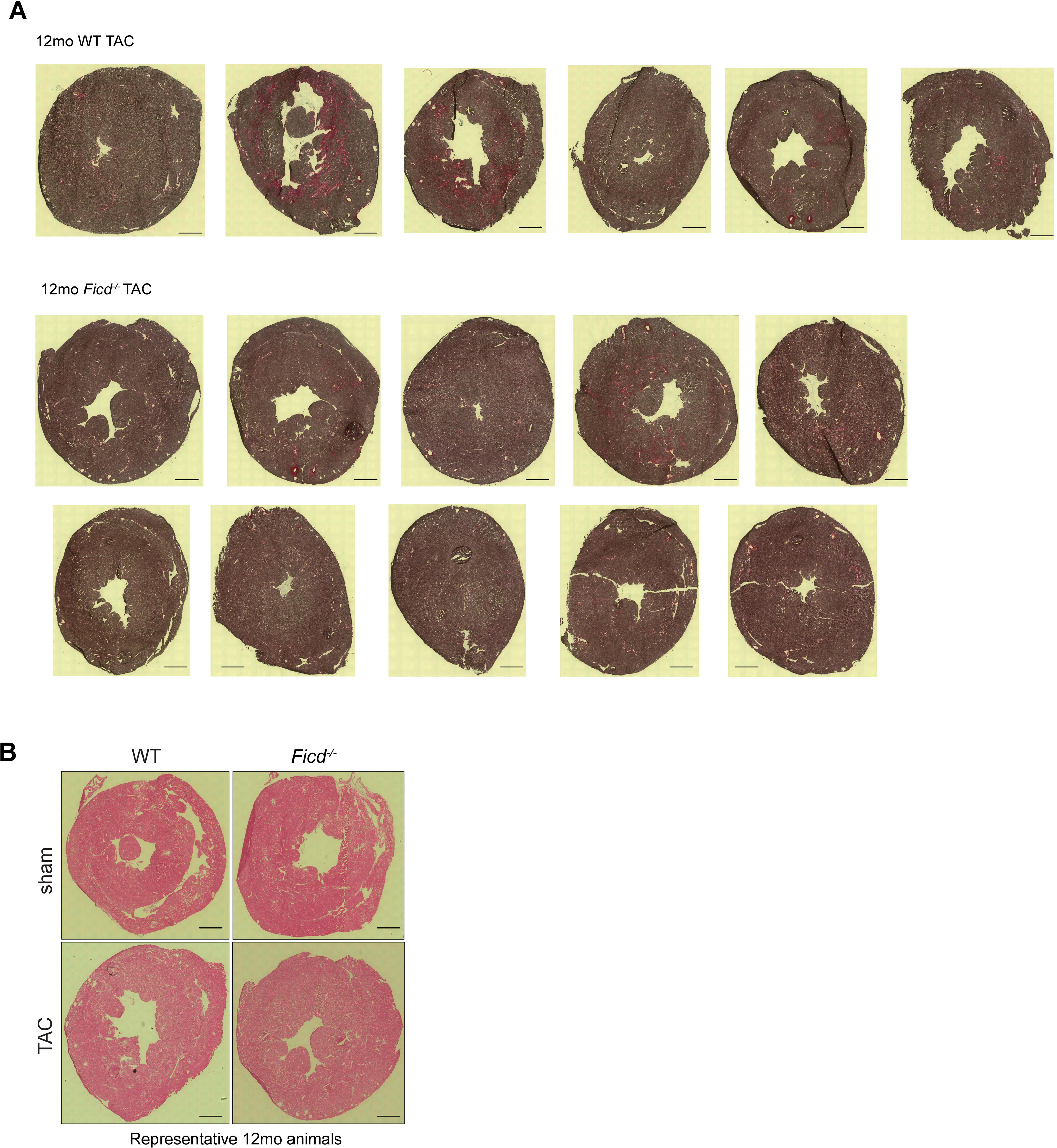
Fibrosis and H&E staining of TAC and sham heart cryosections. Images of collected sham and TAC heart cryosections from 6-month and 12-month animals 8 weeks after respective surgeries and stained for (A) fibrosis using picrosirius stain and (B) cell size using H&E staining (Scale bar, 2mm).

**Supplementary Figure S4.**
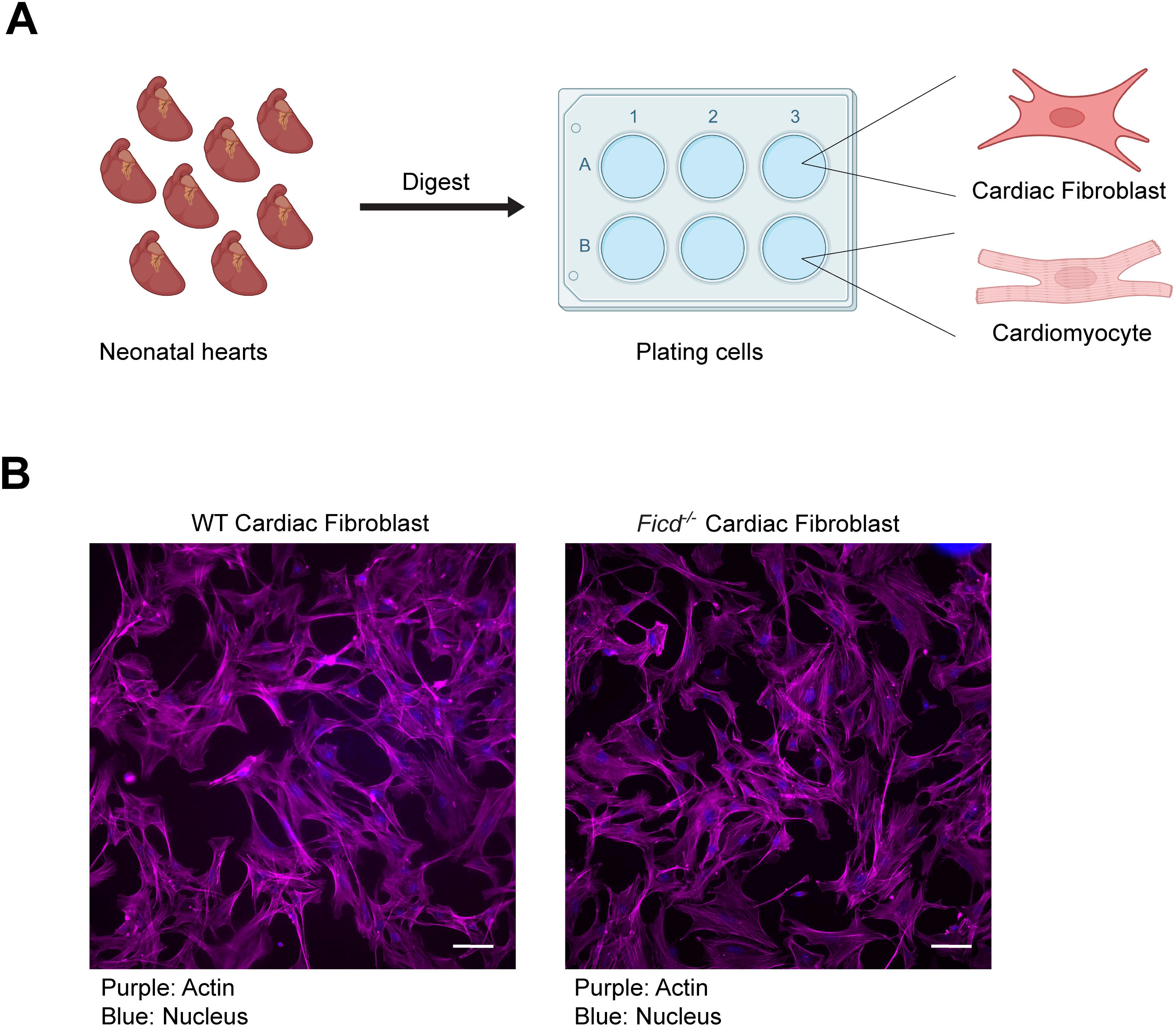
Isolation of primary cardiomyocytes and cardiac fibroblasts. (A) Schematic of isolation protocol. Briefly, the hearts are isolated from p1-3 mice (n: 6-8 mice) and digested overnight. The cells are further digested the following day and cardiomyocytes and cardiac fibroblasts are separated from each other by pre-plating cardiac fibroblasts in uncoated dishes and then washing off and re-plating the cardiomyocytes onto gelatin/collagen pre-coated dishes or 2DMB plates. (B) Representative images of cardiac fibroblasts stained for actin (pink) and nuclei (blue) (Scale bar, 40μm).

**Supplementary Figure S5.**
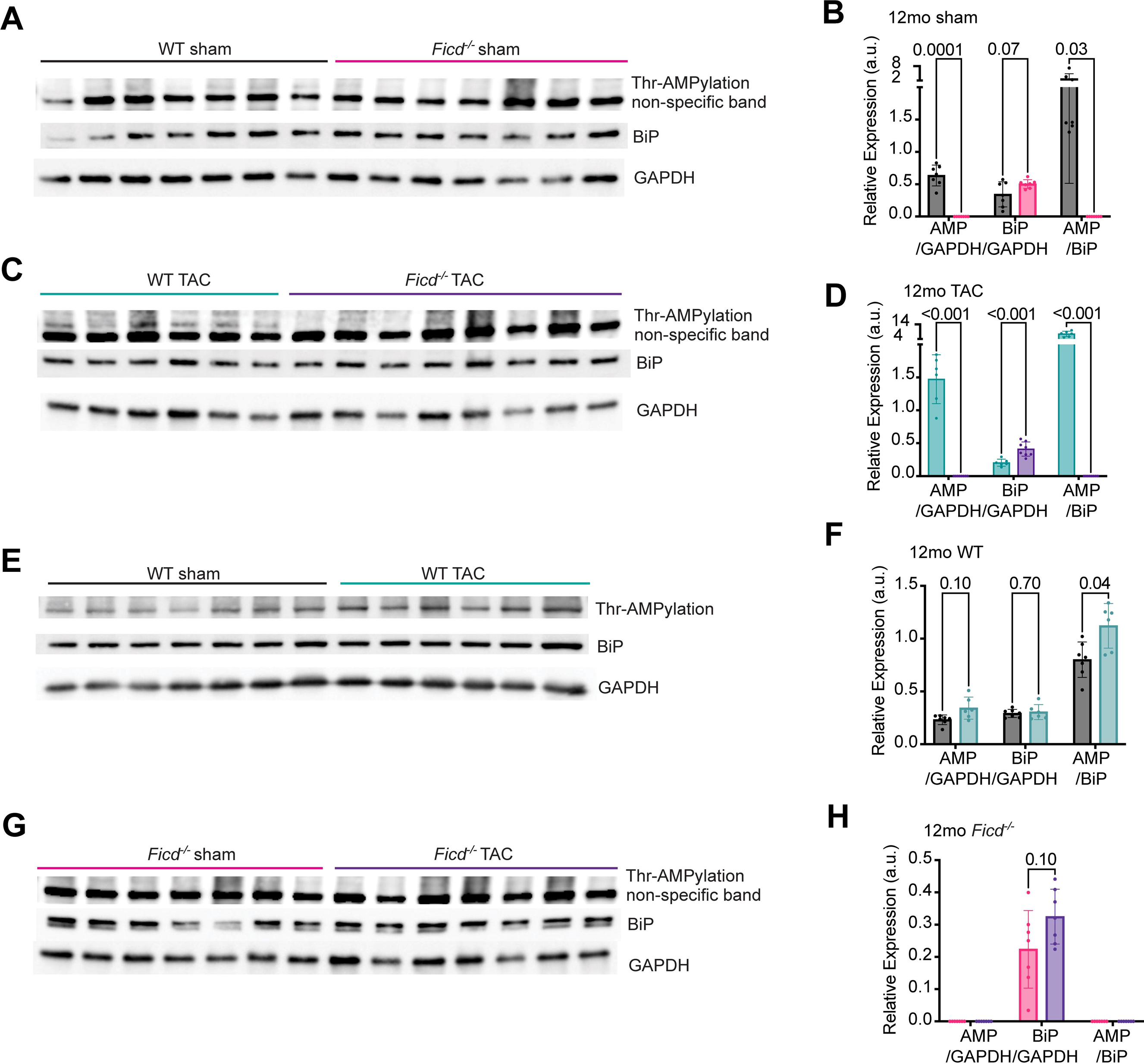
Hypertrophy increases BiP AMPylation in 12-month TAC hearts. Hearts were collected from 14-month mice 8 weeks post-TAC or sham surgery and processed for protein quantification. Protein from (A) sham only, (C) TAC only, (E) WT only, and (G) *Ficd^-/-^*only mice were run on 10% Tris-Glycine SDS acrylamide gels, blocked with 5% BSA, and probed for Thr-AMPylation, BiP, and GAPDH (loading control) levels (n=6-7 mice per genotype per condition). Quantification of Thr-AMPylation, BiP, and GAPDH from (B) sham only, (D) TAC only, (F) WT only, (H) *Ficd^-/-^*only gels. Indicated P values were calculated using unpaired t tests. 2-way ANOVA tests were performed in parallel to confirm significance of results.

